# Universally valid reduction of multiscale stochastic biochemical systems using simple non-elementary propensities

**DOI:** 10.1101/2021.04.08.438974

**Authors:** Yun Min Song, Hyukpyo Hong, Jae Kyoung Kim

## Abstract

Biochemical systems consist of numerous elementary reactions governed by the law of mass action. However, experimentally characterizing all the elementary reactions is nearly impossible. Thus, over a century, their deterministic models that typically contain rapid reversible bindings have been simplified with non-elementary reaction functions (e.g., Michaelis-Menten and Morrison equations). Although the non-elementary reaction functions are derived by applying the quasi-steady-state approximation (QSSA) to deterministic systems, they have also been widely used to derive propensities for stochastic simulations due to computational efficiency and simplicity. However, the validity condition for this heuristic approach has not been identified even for the reversible binding between molecules, such as protein-DNA, enzyme-substrate, and receptor-ligand, which is the basis for living cells. Here, we find that the non-elementary propensities based on the deterministic total QSSA can accurately capture the stochastic dynamics of the reversible binding in general. However, serious errors occur when reactant molecules with similar levels tightly bind, unlike deterministic systems. In that case, the non-elementary propensities distort the stochastic dynamics of a bistable switch in the cell cycle and an oscillator in the circadian clock. Accordingly, we derive alternative non-elementary propensities with the stochastic low-state QSSA, developed in this study. This provides a universally valid framework for simplifying multiscale stochastic biochemical systems with rapid reversible bindings, critical for efficient stochastic simulations of cell signaling and gene regulation. To facilitate the framework, we provide a user-friendly open-source computational package, ASSISTER, that automatically performs the present framework.

## Introduction

To understand the complex dynamics of numerous molecular interactions in living cells, quantitative analysis using mathematical models is essential [1]. While elementary reactions in living cells can be modeled by the law of mass action, characterizing all their kinetics is challenging. Thus, over a century, the combined effect of a set of elementary reactions such as rapid reversible bindings has been described with non-elementary reaction functions (e.g., Michaelis-Menten and Morrison equations) to simplify deterministic models [2–7]. Since the early 2000s, these deterministically driven non-elementary reaction functions have also been widely used to derive propensity functions for stochastic simulations, which greatly reduces the computational cost [8–33]. This heuristic approach for efficient stochastic simulations was believed to be valid as long as the non-elementary reaction functions are accurate in the deterministic sense. However, unfortunately, this was not the case [33–40]. The reason for the discrepancy between the deterministic and stochastic simulations has been recently identified for some cases [37–40], but not for all [41]. Currently, guidelines for this popular but heuristic method for efficient stochastic simulations with non-elementary propensity functions are absent.

The non-elementary reaction functions are mainly the result of the reduction of deterministic models with the following reversible binding reactions:

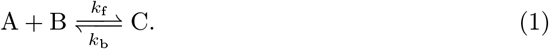

The reversible binding between molecules, such as enzyme-substrate, receptor-ligand, and protein-DNA, is the first step for nearly all biological functions of living cells [42]. However, rather than the reversible binding itself, its outcome is usually our major interest. For instance, rather than the binding between a transcription factor and DNA, we are interested in its outcome, the transcription. Furthermore, the transcription factor binding to DNA takes at most one second while transcription takes about 30 minutes in a mammalian gene [43], which causes stiffness in numerical simulations [44].

Fortunately, such rapid reversible binding reactions can be eliminated from models using the property: the levels of the species (A, B, and C) regulated by the reversible binding more quickly equilibriate to their quasi-steady-states (QSSs) compared with the total levels of the bound and unbound species, which are not affected by the reversible binding. In deterministic models, their quasi-steady-state approximations (QSSAs), which are non-elementary reaction functions, can be obtained by finding the steady-state solution of the associated differential equation in terms of the total variables. Because the QSSAs are determined by the total variables, they are known as the “total” QSSA (tQSSA). After replacing the variables that represent the levels of A, B, and C with their tQSSAs, rapid reversible bindings have been successfully eliminated from various deterministic models describing enzyme catalysis, gene regulation, and cell cycle regulation [5–7, 23, 45–51]. Note that adopting the total variables leads to timescale separation among variables, while the sole rapidity of the reversible binding reactions does not guarantee timescale separation between the original variables, A, B, and C [7] (see Discussion for details).

In stochastic models, the QSSAs for the numbers of A, B, and C are their stationary average numbers (i.e., the first moment) conditioned on the total numbers of the bound and unbound species for uni- or bi-molecular reactions [27–30] (see S1 Appendix for details). These stochastic QSSAs can be obtained by finding the steady-state solution of the chemical master equation (CME). However, unlike the deterministic tQSSA, the stochastic QSSA has a complex form (Eq. (4)), which does not provide any intuition, and importantly, increases computational cost. Thus, its approximation has been derived with the deterministic tQSSA. This approximation, often referred to as the stochastic tQSSA (stQSSA) [7, 31, 38, 39], leads to non-elementary propensity functions for stochastic simulations using the Gillespie algorithm [52]. In this way, the stochastic dynamics of various systems have been accurately captured with low computational cost [7, 30–33, 38, 39, 53, 54]. However, a recent study reported that the stQSSA can be inaccurate [41], which raises the question of validity conditions for the stQSSA.

Here, we identify the complete validity condition for using the stQSSA to simplify stochastic models containing rapid reversible bindings. Specifically, we find that the stQSSA is accurate for a wide range of conditions. However, when two species whose molar ratio is ~1:1 tightly bind, the stQSSA highly overestimates the number of unbound species. In this case, using the stQSSA to simplify stochastic models distorts the stochastic dynamics of the transcriptional repression, the transcriptional negative feedback loop of the circadian clock, and the bistable switch for mitosis. Importantly, by using the fact that the number of the unbound species is low due to the tight binding when the stQSSA is inaccurate, we develop an alternative approach, stochastic “low-state” QSSA (slQSSA). In this way, when reversible bindings are tight and not tight, slQSSA and stQSSA can be used, respectively, which enables one to obtain accurately reduced stochastic models for any case. This proposes a complete and straightforward strategy for efficiently simulating multiscale stochastic biochemical systems containing the fundamental elementary reaction, i.e., rapid reversible binding. To facilitate this framework, we provide a user-friendly open-source computational package, ASSISTER (Adaptive Simplification of StochastIc SystEm with Reversible binding).

## Results

### stQSSA can overestimate the number of the unbound species

In the reversible binding reaction (Eq. (1)), the concentration of A, denoted by 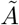, is governed by the following ODE:

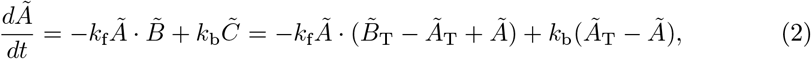

Where 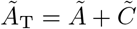 and 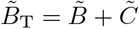 are the total concentrations of the bound and unbound species. By solving 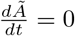 in terms of 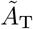 and 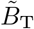, the tQSSA for 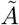 can be obtained as follows:

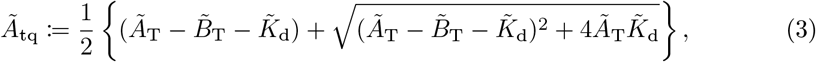

where the 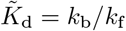 is the dissociation constant. Note that if the reversible binding (Eq. (1)) is embedded in a larger system, there could be other reactions affecting the dynamics of 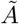 and thus additional terms in Eq. (2). However, as long as the reversible binding is fast (i.e., *k*_f_ and *k*_b_ are much larger than the other reaction rates), 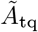 is still an accurate tQSSA for 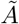. Similarly, by solving 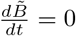 and 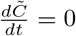, the tQSSAs for 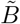 and 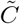 can be obtained. These tQSSAs, also known as the Morrison equations [6], are generally valid, unlike the Michaelis-Menten type equations which are valid only when the enzyme concentration is negligible [7, 47, 48, 50]. Thus, the tQSSAs have been used to simplify models containing not only interactions between metabolites but also proteins whose concentrations are typically comparable [7].

Unlike the deterministic QSSA (Eq. (3)), the stochastic QSSA, which is the stationary average number conditioned on the total numbers of the bound and unbound species, has a complex form [41, 55, 56]. For instance, the stochastic QSSA for the number of A (⟨*A*⟩) can be expressed in terms of the dimensionless variables and parameters, 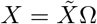, where Ω is the volume of a system (e.g., 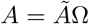, 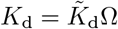 as follows (see Methods for details):

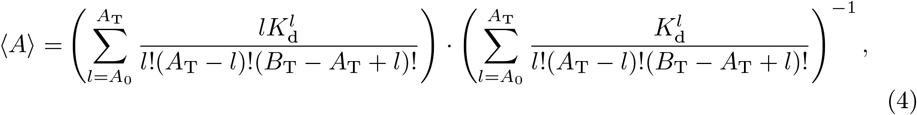

where *A*_0_ = max {0*, A*_T_ − *B*_T_}. This complex form of the stochastic QSSA does not provide any intuition and importantly increases computational cost. Thus, as an alternative to the stochastic QSSA, its approximation, the stQSSA was derived with the deterministic tQSSA [7, 22–26, 31]. Specifically, the stQSSA for *A* (*A*_tq_) can be derived from 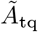 (Eq. (3)) after replacing the concentration-based variables and parameters 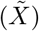 with dimensionless variables and parameters (*X*) as follows:

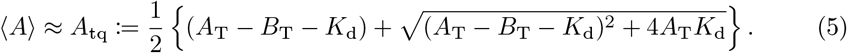

Similarly, the stQSSA for *B* and *C* (*B*_tq_ and *C*_tq_) can be obtained from their deterministic tQSSAs as follows:

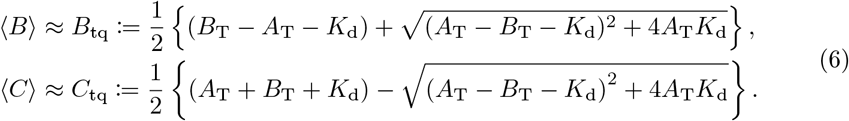

To identify the validity conditions for these stQSSAs, we calculated the relative error (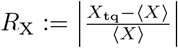, X = A, B, C) of the stQSSA (*X*_tq_) to the stochastic QSSA (⟨*X*⟩) (Fig 1a-1c). The errors are nearly zero in most of the parameter regions, which explains why various stochastic models reduced with the stQSSA have been accurate in most previous studies [7, 30–33, 38, 39, 53, 54]. However, the relative errors of the unbound species (*R*_A_ and *R*_B_) are high when *A*_T_ ≈ *B*_T_. Specifically, the relative error of the bound species (*R*_C_) is at most ~0.2 but that of the unbound species (*R*_A_, *R*_B_) can be ~100.

**Fig 1.**
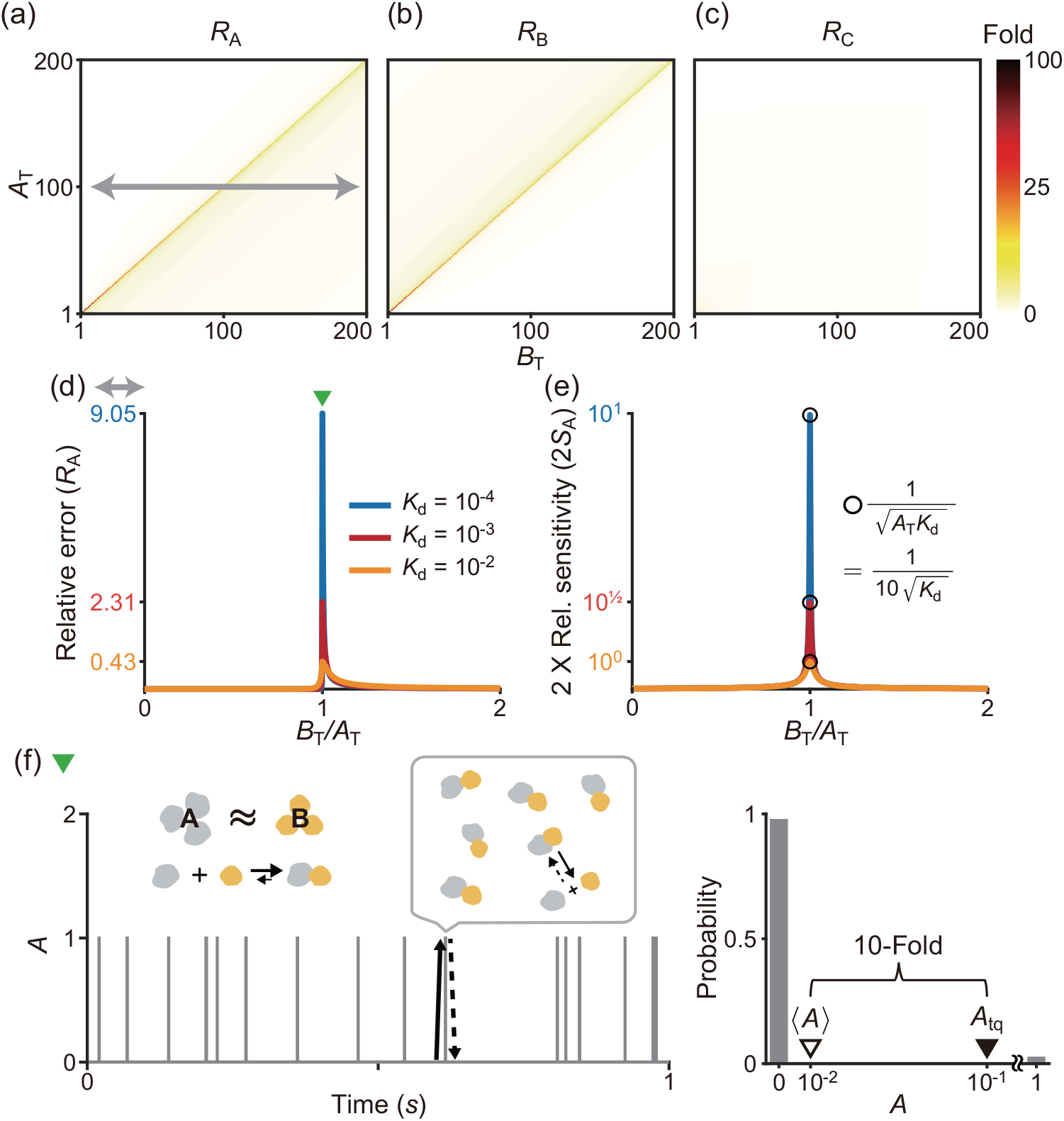
stQSSA overestimates the number of the unbound species when their molar ratio is ~1:1 and binding is tight. **(a-c)** Heat maps of the relative errors (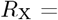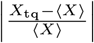) of the stQSSA (*X*_tq_) to the stochastic QSSA (⟨*X*⟩) for *X* = *A, B, C* in the reversible binding reaction (Eq. (1)). Color in the heat maps represents the maximum value of *R*_X_ calculated by varying *K*_d_ from 10^−4^ to 10^2^ for each total number of the bound and unbound species (*A*_T_ = *A* + *C* and *B*_T_ = *B* + *C*). *R*_A_ and *R*_B_ can be extremely large when *A*_T_ ≈ *B*_T_ while *R*_C_ is always small. **(d)** *R*_A_ calculated over *B*_T_*/A*_T_ between 0 and 2 (gray arrow in a) for three fixed *K*_d_ values (10^−4^, 10^−3^ and 10^−2^). *R*_A_ becomes larger as *B*_T_*/A*_T_ is similar to 1 and the *K*_d_ becomes smaller (i.e., the binding becomes tighter). **(e)** *R*_A_ mainly depends on the relative sensitivity of *A*_tq_ (i.e., 2*S*_A_), which can be derived in a simple form, unlike *R*_A_ (Eq. (8)). The maximum value of 2*S*_A_ is given by 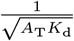, which is achieved when *B*_T_*/A*_T_ is similar to 1 as in the case of *R*_A_. **(f)** A trajectory (left) and the stationary probability distribution (right) of *A* for a parameter set where *R*_A_ is large (green triangle in d, *A*_T_ = *B*_T_ = 100, *k*_f_*/*Ω = 10^4^*s*^−1^*, k*_b_ = 1*s*^−1^), simulated using the Gillespie algorithm. Since *A*_T_ = *B*_T_ and A binds with B tightly, *A* = 0 (i.e., every A is bound) most of the time, and it rarely becomes 1 by the weak unbinding reaction (solid arrow) and immediately comes back to 0 by the strong binding reaction (dotted arrow). As a result, when *K*_d_ = 10^−4^, the probability that *A* = 1 is ~0.01, but the stQSSA for *A* overestimates it as ~0.1, which is 10 times larger (i.e., a 10-fold error)

To investigate why *R*_A_ is high when *A*_T_ ≈ *B*_T_, we derived the exact upper and lower bounds for *R*_A_ (see Methods for details):

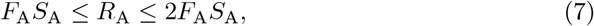

where *F*_A_ is the Fano factor of *A* (i.e., 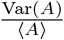), and *S*_A_ is the relative sensitivity of *A*_tq_ with respect to *B*_T_ (i.e., 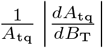). Furthermore, we proved that the Fano factor (*F*_A_) is less than 1 (i.e., *A* has a sub-Poissonian stationary distribution; see S1 Appendix for details). Therefore, *R*_A_, especially its upper bound, mainly depends on *S*_A_ (Figs 1d, 1e, and S1) whose formula can be derived in the following simple form, unlike *R*_A_:

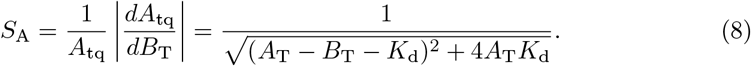

Because *S*_A_ attains the maximum value 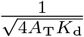 at *B*_T_ = *A*_T_ − *K*_d_, *S*_A_ has a large maximum value when *K*_d_ ≪ 1 at *A*_T_ = *B*_T_ + *K*_d_ ≈ *B*_T_. This explains why *R*_A_, whose upper bound is mainly determined by 2*S*_A_, is large when the binding is tight (*K*_d_ ≪ 1) and the total numbers of the bound and unbound species are similar (*A*_T_ ≈ *B*_T_) (Fig 1d). In this case, the majority of A is bound with B, and thus *A* = 0 most of the time (Fig 1f left). That is, *A* rarely becomes 1 by the weak unbinding reaction and then immediately *A* becomes 0 by the strong binding reaction. As a result, the probability that *A* = 1 is approximately 1% (i.e., ⟨*A*⟩ ≈ 0.01), but the stQSSA for *A* (*A*_tq_) overestimates it as 10%, which is 10 times larger (Fig 1f right). Since A and B are symmetric, the above analysis can be applied to B, analogously.

### stQSSA can overestimate the transcriptional activity

We found that the stQSSA for the number of the unbound species is inaccurate if their molar ratio is ~1:1 and their binding is tight (Fig 1d-1f). Thus, we expected that in such cases, using the stQSSA to eliminate a rapid reversible binding in a stochastic model can distort its dynamics. To illustrate this, we constructed a simple gene regulatory network where gene expressions are determined by a reversible binding between transcription factors and genes (Fig 2a left, Table S1); DNA (D) and a transcription factor (P) reversibly bind to form a complex (D:P). As P acts as a repressor of M_R_ transcription, the transcription rate of M_R_ is proportional to the number of the unbound DNA (*D*). On the other hand, as P acts as an activator of M_A_ transcription, the transcription rate of M_A_ is proportional to the number of the bound DNA (*D*:*P*). Note that the number of unbound and bound DNA can be interpreted as the number of unbound and bound DNA binding sites. In this model, because the reversible binding reaction between D and P is much faster than the other reactions (i.e., the production and the decay of M_R_ and M_A_), the variables (*D* and *D*:*P*) rapidly reach their QSS. Thus, by replacing them with their stQSSAs (*D*_tq_ and *D*:*P*_tq_), we can obtain a reduced model (Fig 2a right, Table S2). The reduced model consists of only the slow variables, *M*_R_ and *M*_A_, because *D*_tq_ and *D*:*P*_tq_ are fully determined by the conserved total number of the DNA (*D*_T_ = *D* + *D*:*P*) and the conserved total number of the transcription factor (*P*_T_ = *P* + *D*:*P*), as illustrated in Table S2. This elimination of the fast variables, which are the major source of computational cost, greatly reduces the computation time of stochastic simulations [27–29].

**Fig 2.**
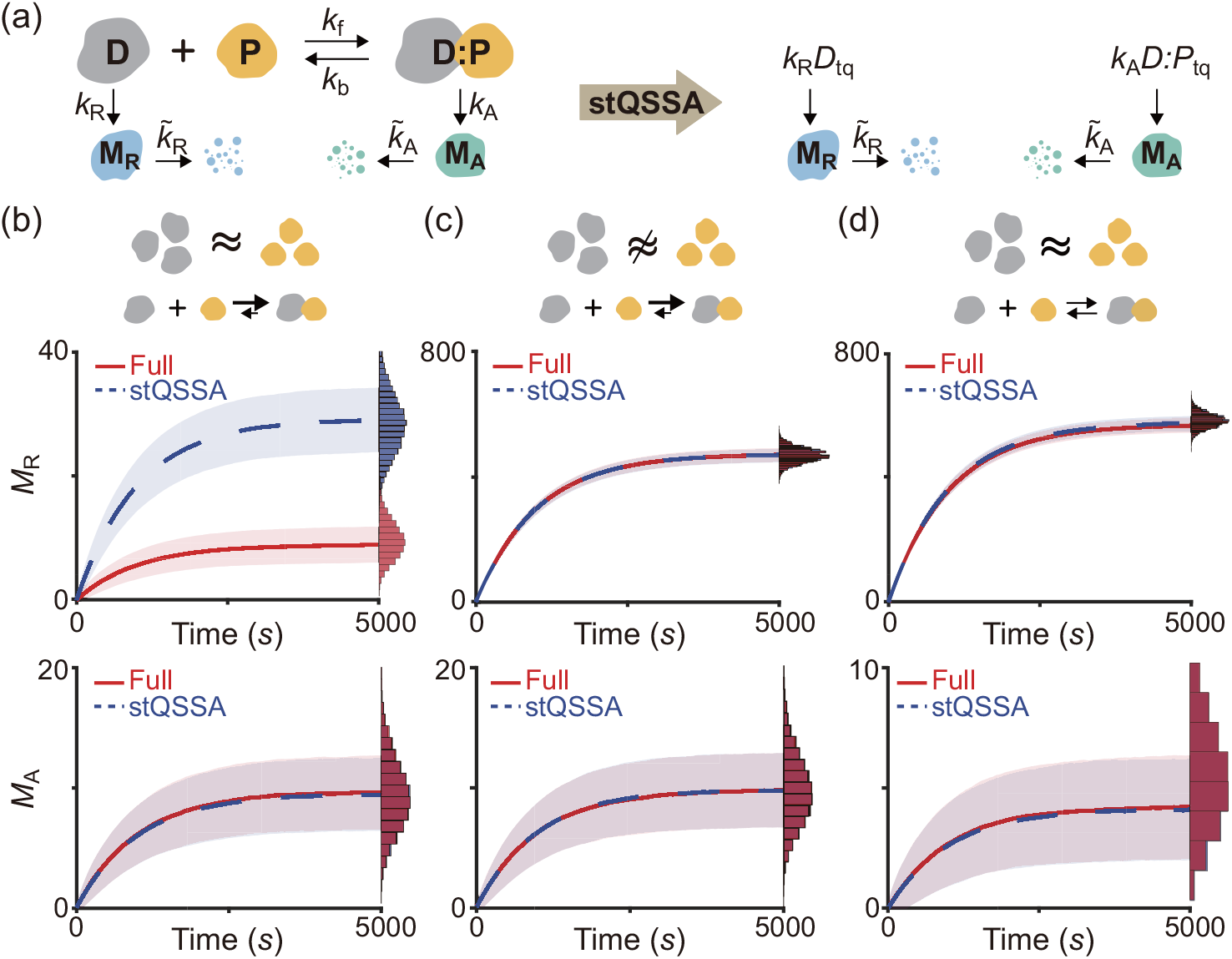
When DNA and a transcription factor bind tightly and their levels are similar, the stQSSA overestimates the number of the unbound DNA. **(a)** Full model diagram of a gene regulatory network containing a rapid reversible binding between DNA (D) and a transcription factor (P) to form a complex (D:P) (left, Table S1). The transcription rates of M_R_ and M_A_ are proportional to *D* and *D*:*P*, respectively. By replacing *D* and *D*:*P* with their stQSSAs (*D*_tq_ and *D*:*P*_tq_), we can obtain a reduced model which consists of only slowly varing *M*_R_ and *M*_A_ (right, Table S2). **(b-d)** Trajectories of *M*_R_ (top) and *M*_A_ (bottom) from the full model (red) and the reduced model (blue) simulated using the Gillespie algorithm (see Tables S1 and S2 for propensity functions). The lines with colored ranges and the histograms represent the mean standard deviation and the stationary distribution of 10^4^ trajectories, respectively. When *D*_T_ and *P*_T_ are the same (*D*_T_ = *P*_T_ = 10) and the binding is tight (*K*_d_ = 10^−2^), the *M*_R_ trajectories simulated with the reduced model largely exceed those simulated with the full model (b top) because *D*_tq_ overestimates the stochastic QSSA for *D* (⟨*D*⟩). On the other hand, *D*:*P*_tq_ accurately approximates the stochastic QSSA for *D*:*P* (⟨*D*:*P*⟩), and thus the reduced model accurately captures the dynamics of *M*_A_ (b bottom). If *D*_T_ is not similar to *P*_T_ (*D*_T_ = 15*, P*_T_ = 10) (c) or the binding is weak (*K*_d_ = 10) (d), *D*_tq_ and *D*:*P*_tq_ accurately approximate ⟨*D*⟩ and ⟨*D*:*P*⟩, respectively, so that the reduced model accurately captures the dynamics of both *M*_R_ and *M*_A_ of the full model.

To test whether the reduced model accurately captures the dynamics of the full model, we compared their stochastic simulations with the Gillespie algorithm (see Tables S1 and S2 for propensity functions) [52]. When *D*_T_ and *P*_T_ are the same and the binding between D and P is tight, *M*_R_ simulated with the reduced model largely exceeds *M*_R_ simulated with the full model (Fig 2b top) because the stQSSA (*D*_tq_) overestimates the stochastic QSSA for the number of the unbound DNA (⟨*D*⟩) which determines the transcription rate of M_R_ (Fig 2a), as seen in Fig 1f. On the other hand, when *D*_T_ is not similar to *P*_T_ (Fig 2c top) or the binding is weak (Fig 2d top), *D*_tq_ accurately approximates ⟨*D*⟩ as seen in Fig 1d, and thus the reduced model accurately captures the dynamics of *M*_R_ in the full model.

Unlike *M*_R_ (Fig 2b top), the stochastic dynamics of *M*_A_ of the reduced model and the full model are identical (Fig 2b-2d bottom) because the stQSSA for *D*:*P* (*D*:*P*_tq_) always accurately approximates the stochastic QSSA for the number of the bound DNA (⟨*D*:*P*⟩) which determines the transcription of *M*_A_ (Fig 1c). Taken together, the stQSSA can be used to describe transcriptional activation depending on bound DNA under any conditions (Fig 2b-2d bottom). On the other hand, it needs to be restrictively used to describe transcriptional repression depending on unbound DNA (Fig 2b-2d top).

### stQSSA can distort oscillatory dynamics

To illustrate how the stQSSA distorts the dynamics when the molar ratio between tightly binding species is ~1:1, we investigated the simple model where the molar ratio is conserved (Fig 2). However, the total copy numbers of binding species and thus their molar ratio can be varied (e.g., oscillate) in a living cell due to other reactions in a larger system. This raises the question of whether the model reduction based on the stQSSA is accurate or not if the molar ratio is temporarily ~1:1. To investigate this, we used a modified Kim-Forger model, which describes the transcriptional negative feedback loop of the mammalian circadian clock [24, 49, 51]. In this model (Fig 3a top, Table S3), free activator (A) promotes the transcription of mRNA (M), and the protein translated from M produces repressor (R) passing through several steps (P_*i*_, *i* = 1, 2, 3). Then R reversibly binds with A to form a complex (A:R) which no longer promotes the transcription, and thus represses its own transcription. In this model, the reversible binding between R and A is much faster than the other reactions (i.e., production and decay). Thus, by replacing the fast variable *A*, which determines the transcription rate of M, with its stQSSA (*A*_tq_), we can obtain a reduced model (Fig 3a bottom, Table S4). The reduced model consists of only the slow variables, *R*_T_, *M* and *P_i_*, because *A*_tq_ is fully determined by the conserved total number of the activator (*A*_T_ = *A* + *A*:*R*) and the slowly varying total number of the repressor (*R*_T_ = *R* + *A*:*R*), as illustrated in Table S4.

**Fig 3.**
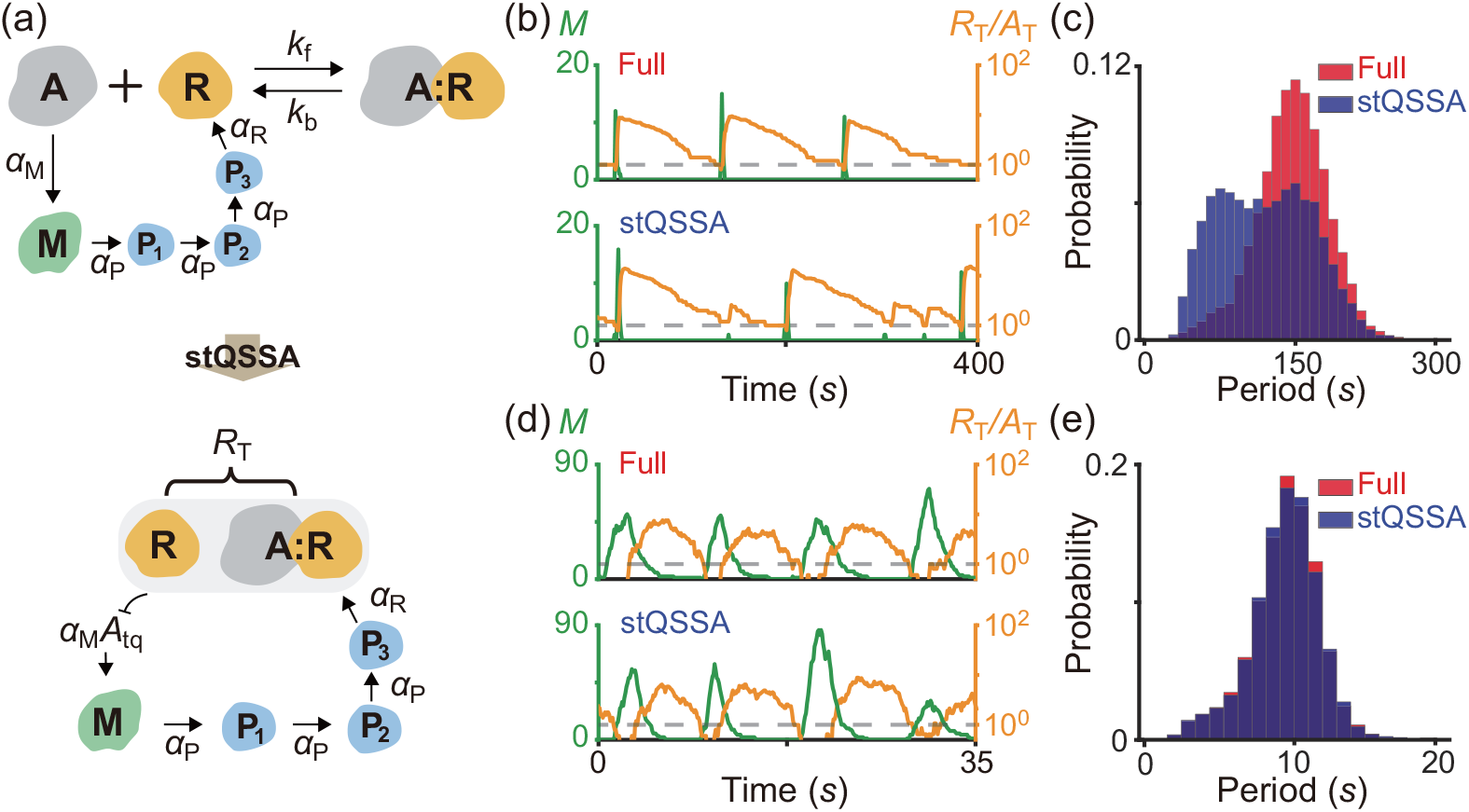
stQSSA can distort the dynamics of a biological oscillator. **(a)** Full model diagram of an oscillatory transcriptional negative feedback loop (top, Table S3). Unbound activator (A) promotes the transcription of mRNA (M), and the protein translated from M produces repressor (R) passing through several steps (P_*i*_, *i* = 1, 2, 3). Then R binds with A to form a complex (A:R) which is transcriptionally inactive, and thus represses its own transcription. As the reversible binding between R and A is rapid, by replacing *A* with its stQSSA (*A*_tq_), we can obtain a reduced model which consists of only slowly varying *R*_T_, *M*, and *P_i_* (bottom, Table S4). **(b-c)** Oscillatory trajectories of *M* (green) and *R*_T_*/A*_T_ (orange) simulated with the full model (b top) and the reduced model (b bottom), using the Gillespie algorithm (see Tables S3 and S4 for propensity functions). When R binds with A tightly (*K*_d_ = 10^−4^) both the full model and the reduced model show the oscillatory behaviors. However, when the trajectory of *R*_T_*/A*_T_ stays near 1 (dashed lines in b), *A*_tq_ overestimates the stochastic QSSA for *A* (⟨*A*⟩), and thus the transcription more frequently occurs in the reduced model (b bottom) compared to the full model (b top). As a result, the reduced model predicts a shorter period than the full model (c). **(d-e)** On the other hand, when the degradation rate of R increases and thus the trajectory of *R*_T_*/A*_T_ stays near 1 for a short time (d; dashed lines), the reduced model accurately captures the dynamics of the full model (e).

In the model, because R tightly binds with A, when *R*_T_*/A*_T_ ≈ 1, *A*_tq_ overestimates the stochastic QSSA for *A* (⟨*A*⟩) and thus the transcription rate of M. As a result, when the trajectory of *R*_T_*/A*_T_ reaches close to 1 (dashed lines in Fig 3b), the transcription more frequently occurs in the reduced model (Fig 3b bottom) compared to the full model (Fig 3b top). This overestimated transcriptional activity leads to the shorter peak-to-peak periods of the reduced model compared to the full model (Fig 3c). On the other hand, when the degradation rate of R increases and thus the trajectory of *R*_T_*/A*_T_ stays near 1 for an extremely short time (Fig 3d dashed lines), the reduced model accurately captures the dynamics of the full model (Fig 3e). Taken together, if ~1:1 molar ratio between the tightly binding activator and repressor of the transcriptional negative feedback loop persists for a considerable time, using the stQSSA overestimates the transitional activity and thus the frequency of oscillation.

### stQSSA can distort bistable dynamics

To investigate how the misuse of the stQSSA distorts the dynamics of a bistable switch, we used a previously developed bistable switch model for the maturation promoting factor, cyclin B/Cdc2, whose activation promotes mitosis (Fig 4a top, Table S5) [46, 57]. In the model, the inactive form of cyclin B/Cdc2 (P) is converted to an active form (M) by Cdc25 (D). Furthermore, as M activates D which converts P to M, M promotes its own activation (i.e., form a positive feedback loop; see [46, 57] for details). The positive feedback loop is suppressed by Suc1 protein (B) as it binds with M to form a complex (M:B) which no longer activates D. The total activated cyclin B/Cdc2 (M and M:B) become P with the same constant rate. In this model, the reversible binding between M and B is much faster than the other reactions. Thus, by replacing the fast variable *M* with its stQSSA (*M*_tq_) a reduced model can be derived (Fig 4a bottom, Table S6). The reduced model consists of only the slow variables, *M*_T_ and *P*, because *M*_tq_ is fully determined by the conserved total number of Suc1 (*B*_T_ = *B* + *M*:*B*) and the slowly varying total number of the activated cyclin B/Cdc2 (*M*_T_ = *M* + *M*:*B*), as illustrated in Table S6.

**Fig 4.**
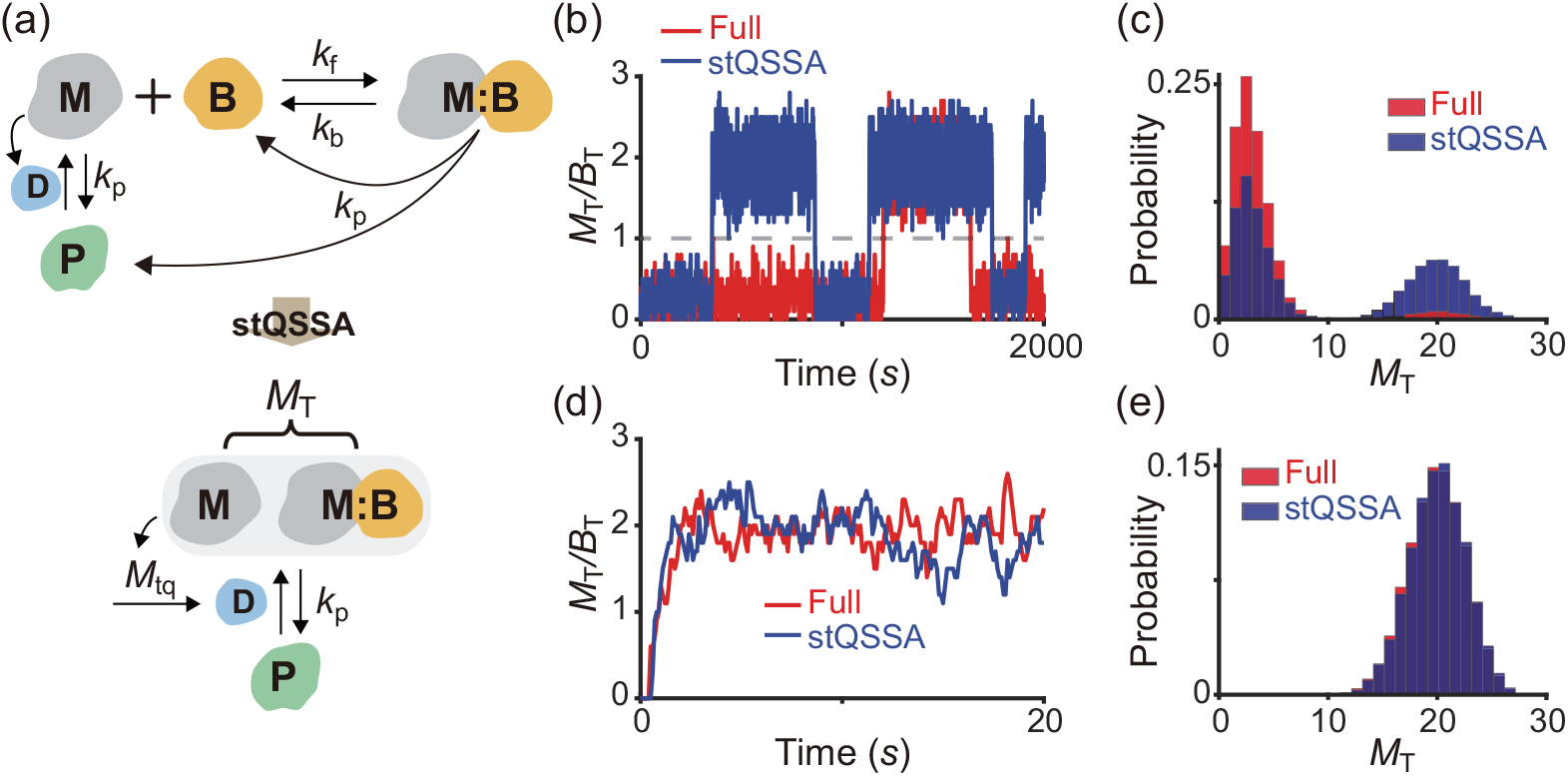
stQSSA can distort the dynamics of a bistable switch. **(a)** Full model diagram of a bistable switch for mitosis (top, Table S5). The inactive form of cyclin B/Cdc2 (P) becomes an active form (M) by Cdc25 (D). In this process, M enhances its own activation by activating D, and thus forms a positive feedback loop (see [46, 57] for details). The positive feedback loop is suppressed as Suc1 protein (B) binds with M to form a complex (M:B) which does not activates D. The total activated cyclin B/Cdc2, M and M:B, becomes P with the same constant rate. As the reversible binding between M and B is rapid, by replacing *M* with its stQSSA (*M*_tq_), we can obtain a reduced model which consists of only slowly varying *M*_T_ and *P* (bottom, Table S6). **(b-c)** Simulated trajectories (b) and the stationary distributions (c) of *M*_T_ from the full model and the reduced model using the Gillespie algorithm (see Tables S5 and S6 for propensity functions). When M binds with B tightly (*K*_d_ = 10^−3^), both the full model and the reduced model show the bistable behaviors between the upper and lower modes, which are separated by *M*_T_*/B*_T_ = 1 (dashed line in b). However, because *M*_tq_ overestimates the stochastic QSSA for *M* (⟨*M*⟩) when *M*_T_*/B*_T_ is close to 1, the trajectory from the reduced model is more attracted to the upper mode compared to the full model (b). As a result, the bimodal distribution of *M*_T_ from the reduced model is biased to the upper mode (c). **(d-e)** On the other hand, when the binding between M and B becomes weak (*K*_d_ = 10), *M*_tq_ accurately estimates ⟨*M*⟩, and thus the reduced model accurately captures the dynamics of the full model, which no longer shows the bistable behavior.

When M and B tightly bind, both the full model and the reduced model show the bistable behaviors (i.e., bimodal stationary distributions) of *M*_T_ (Fig 4b). However, the trajectory of the reduced model is more attracted to the upper mode of *M*_T_ compared to the full model (Fig 4b and 4c). This dynamics biased to the upper mode occurs because *M*_tq_ overestimates the stochastic QSSA for *M* (⟨*M*⟩) near the *M*_T_*/B*_T_ = 1 region (Fig 4b dashed line) which separates the upper and lower modes. On the other hand, when the binding between M and B becomes weak, *M*_tq_ accurately approximates the stochastic QSSA for *M* even when *M*_T_ is similar to *B*_T_. Thus, the reduced model accurately captures the dynamics of the full model, which no longer shows bistable behavior (Fig 4d and 4e). Taken together, when the binding between activated Cyclin B/Cdc2 and Suc1 protein is tight, which is essential to generate the bistable switch, using the stQSSA overestimates the activation of Cyclin B/Cdc2 and distorts the dynamics of the bistable switch.

### An alternative approach when the stQSSA is not applicable

In the presence of a rapid and tight reversible binding between species whose molar ratio is ~1:1, the reduction of stochastic models with the stQSSA for the number of the unbound species can cause errors (Figs 2b, 3c, and 4c). In such cases, due to the tight binding, the two species tend to bind until no molecules of one species are left (Fig 1f). Specifically, if *A*_T_ ≤ *B*_T_ (*A*_T_ ≥ *B*_T_), the majority of the A (B) will be bound. Thus, in the presence of tight binding, we can assume that the stationary distributions of *A* or *B* are concentrated on 0 and 1. This low-state assumption allows us to derive the simple approximation for the stochastic QSSA (⟨*A*⟩ in Eq. (4)) (see Methods for details):

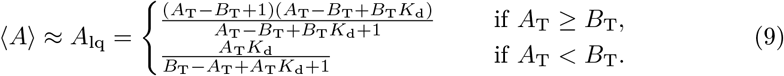

We will refer to this approximation as the stochastic “low-state” QSSA (slQSSA). Since this approximation relies on the property that the state space is restricted at the low level, the stochastic QSSA with singular perturbation analysis introduced in [58] could be used to derive an alternative approximation.

The accuracy of the slQSSA for *A* (Eq. (9)) is expected to increase when *A*_T_*K*_d_ decreases because *A*_T_*K*_d_ is an approximated number of the unbound A. On the other hand, the accuracy of the stQSSA for *A* decreases as *A*_T_*K*_d_ decreases (Fig 1d). To investigate this, we calculated the maximum relative error of 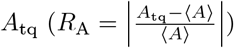 and 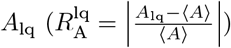 to the stochastic QSSA for *A* (⟨*A*⟩) for each *A*_T_*K*_d_ and *K*_d_ (Fig 5a and 5b). As expected, when *A*_T_*K*_d_ is low and high, the slQSSA and the stQSSA are accurate, respectively. In particular, when *A*_T_*K*_d_ < 10^−1^ and *A*_T_*K*_d_ > 10^1^, 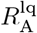 and *R*_A_ are less than 0.1 (i.e., the relative errors are less than 10%), respectively.

**Fig 5.**
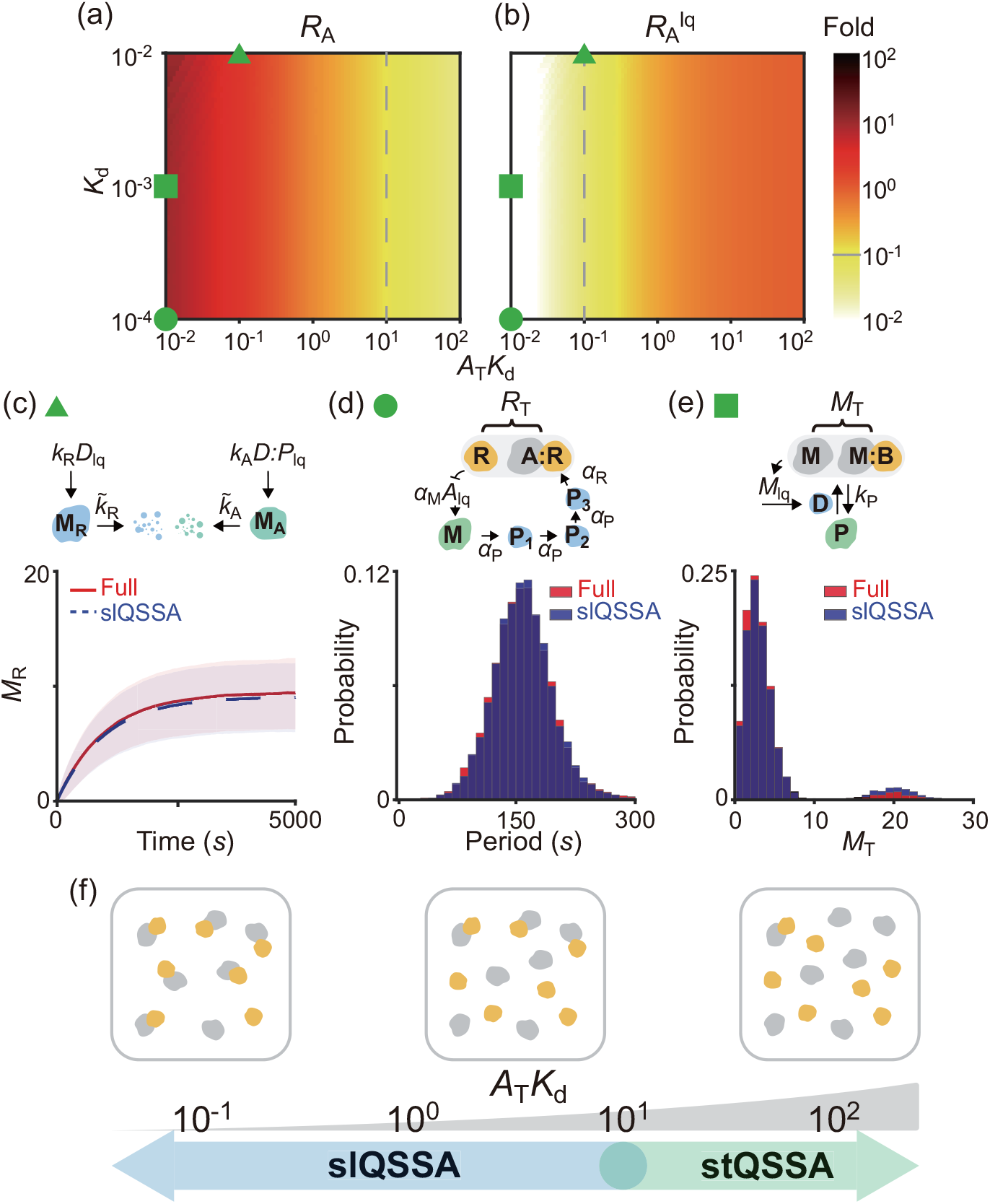
slQSSA can be used to reduce multiscale stochastic biochemical systems containing rapid reversible bindings when the stQSSA is not applicable. **(a-b)** Heat maps of the relative errors 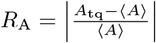 and 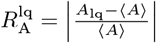 when the stQSSA (*A*_tq_) and the two-state slQSSA (*A*_lq_) approximate the stochastic QSSA for *A* (⟨*A*⟩) in the reversible binding reaction (Eq. (1)). Color represents the maximum value of *R*_A_ and 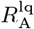 for each *A*_T_*K*_d_ and *K*_d_ when *B*_T_ varies, and the dashed lines represent when those values are 0.1. When *A*_T_*K*_d_ are high and low, the stQSSA and the slQSSA are accurate, respectively. The parameters used in Figs 2b (triangle), 3b (circle), and 4b (square) are located in the region where the slQSSA (b), but not the stQSSA (a), is accurate (the circle is actually located outside of the heat maps; *A*_T_*K*_d_ = 5 × 10^−4^ and *K*_d_ = 10^−4^). **(c-e)** As a result, the full models are successfully reduced with the slQSSA (c-e) but not the stQSSA (Figs 2b, 3c, and 4c). See Tables S2, S4, and S6 for the propensity functions used for the simulations and Fig S3 for a benchmark comparison with GillesPy2, one of the major, standard software suites for stochastic simulation [59]. **(f)** The adaptive use of the stQSSA and the slQSSA to approximate the stochastic QSSA for *A* when *A*_T_*K*_d_ > 10^1^ and otherwise, respectively, guarantees the successful reduction of stochastic models containing rapid reversible bindings. Note that when 10^−1^ < *A*_T_*K*_d_ < 10^1^, the slQSSAs with more than two states need to be used (see Fig S2 for details).

The parameters used in Figs 2b (triangle), 3b (circle), and 4b (square) are located in the region where the stQSSA is inaccurate (Fig 5a) but the slQSSA is accurate (Fig 5b). Therefore, with these parameters, the reduced models obtained by using the slQSSAs accurately capture the dynamics of the full models for the simple gene regulatory network (Fig 5c, Table S2), the transcriptional negative feedback loop (Fig 5d, Table S4), and the bistable switch for mitosis (Fig 5e, Table S6), unlike the stQSSA (Figs 2b, 3c, and 4c). Furthermore, by allowing *A* or *B* to reach more than two states (e.g., 0, 1, and 2), more accurate slQSSAs can be derived (see Methods for details). In particular, the relative errors of the slQSSAs derived by allowing the 3/4/5 states are less than 0.1 when *A*_T_*K*_d_ is less than 2/5/10, respectively (Fig S2). Consequently, for the error tolerance of 0.1, if *A*_T_*K*_d_ < 10^1^ and thus the stQSSA is inaccurate, the slQSSA can be used to approximate the stochastic QSSA for *A* (Fig 5f). Taken together, by using either stQSSA or slQSSA depending on *A*_T_*K*_d_, we can always accurately reduce multiscale stochastic biochemical systems with rapid reversible bindings. Of course, for a different error tolerance, we need a different threshold of *A*_T_*K*_d_ and the number of states for the slQSSA. To facilitate the calculation of such change depending on the error tolerance, we have developed a user-friendly open-source computational package, ASSISTER (Fig 6). In particular, the function `Gillespie_Reduction` in the package automatically constructs a reduced model adaptively using the more accurate one between the two approximation methods and performs accurate and efficient stochastic simulations (see S1 Appendix for the manual).

**Fig 6.**
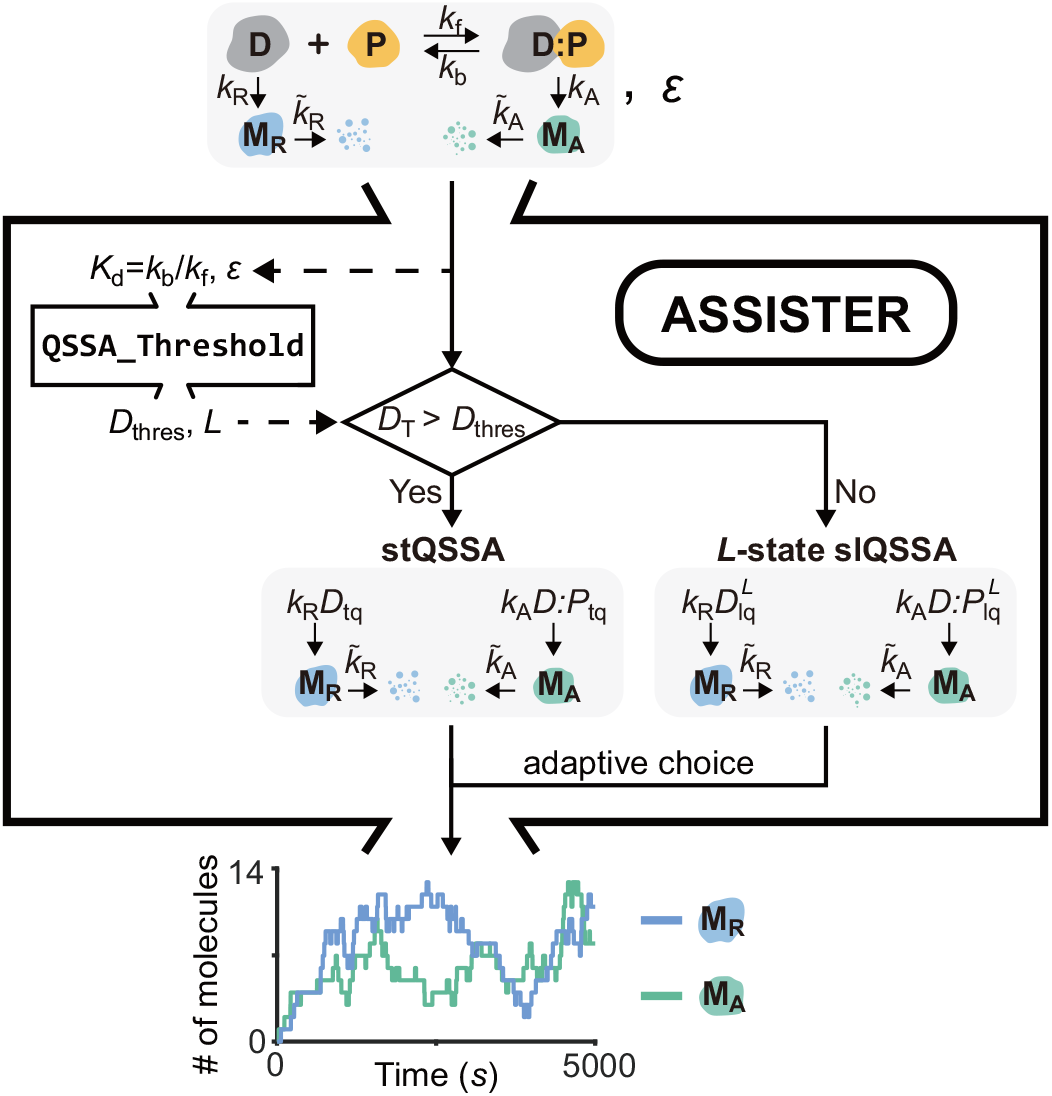
Schematic diagram for the computational package, ASSISTER. When a stochastic model containing a rapid reversible binding 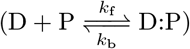 and the error tolerance (*ϵ*) are given as inputs, the auxiliary function `QSSA_Threshold` determines the threshold of the total number of binding molecules, *D*_thres_, and the number of states for the slQSSA, *L*. When the *D*_T_ = *D* + *D*:*P* is less (larger) than *D*_thres_, the *L*-state slQSSA (stQSSA) approximates the exact stochastic QSSA with a relative error less than *ϵ*. Based on the relationship between *D*_thres_ and *D*_T_, the more accurate one between the two models is adaptively chosen. In this way, `Gillespie_Reduction` performs efficient and accurate stochastic simulations, yielding the simulated trajectories as the final output. See the manual in S1 Appendix for a more detailed description of the input and output.

## Discussion

Reversible binding between molecules—for example, between DNA and a transcription factor, a ligand and a receptor, and an enzyme and a substrate—is a fundamental reaction for numerous biological functions [42]. As the reversible binding reactions occur typically on a timescale of 1~1000ms, which is much faster than the other reactions (e.g., 30min for a mammalian mRNA transcription or a protein translation and 10h for their typical lifetimes) [43], a system containing the rapid reversible binding becomes a multi-timescale system. In such multi-timescale systems, the rapid reversible binding prohibitively increases the computational cost of stochastic simulations. Accordingly, to accelerate stochastic simulations, various methods have been developed [44, 60]. In particular, the model reduction using the stQSSA has successfully simplified various stochastic models in numerous studies [7, 30–33, 38, 39]. Thus, it has been commonly believed that the stQSSA is generally accurate for any conditions, until a recent counterexample was identified [41]. In this work, we rigorously derived the validity conditions for using the stQSSA to reduce stochastic models with a rapid reversible binding. Specifically, we showed that the relative error of the stQSSA for the number of unbound species (*R*_A_) mainly depends on the relative sensitivity of the stQSSA (*S*_A_, Eq. (8)), which attains maximum value 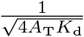 at *A*_T_ = *B*_T_ + *K*_d_. This allowed us to find that the stQSSA for the number of the unbound species is inaccurate if their molar ratio is ~1:1 and their binding is tight (Fig 1f). In that case, the stQSSA highly overestimates the number of the unbound species. Therefore, the reduced models obtained by using the stQSSA distort the dynamics of the gene regulatory model (Fig 2b), the transcriptional negative feedback loop model for circadian rhythms (Fig 3c), and the bistable switch model for mitosis (Fig 4c).

When the reversible binding reactions are sufficiently faster than the other reactions, the deterministic tQSSA is known to be accurate [7, 48, 50]. Indeed, for all examples considered in our work (Figs 2b, 3b, and 4b), the deterministic simulations with the tQSSA are accurate, unlike the stochastic simulations. This indicates that it is risky to investigate the validity conditions of the stQSSA solely based on the validity conditions of the deterministic tQSSA. Instead, the direct derivation of the relative error of the stQSSA is needed, as demonstrated in this study (Eq. (7)). It would be interesting in future work to perform such error analysis for more complex examples, such as coupled enzymatic networks with multiple rapid reversible bindings [26, 61, 62].

The rapid reversible reactions are typical conditions for model reductions using the “partial-equilibrium approximation,” which confines species concentrations to the equilibrium states. In general, this condition does not imply a timescale separation between the variables (*A*, *B*, and *C*), limiting the application of the QSSA. However, the rapid reversible binding guarantees that the total variables *A*_T_ and *B*_T_ always evolve more slowly than the variables *A*, *B*, and *C*. Therefore, the QSSAs in terms of the total variables can lead to accurate model reductions in both deterministic [7, 47, 48, 50] and stochastic [27–30] regimes. In this work, we investigated under which conditions the complex stochastic QSSA (Eq. (4)) can be approximated by the corresponding simple deterministic QSSA (Eq. (5)), referred to as the stQSSA. We found that such an approximation becomes inaccurate when two species with similar levels tightly bind. Thus, we derived an alternative approximation for the stochastic QSSA: slQSSA. Using the two approximations for the stochastic QSSA, we developed the universally valid reduction framework for stochastic models containing rapid reversible bindings. We also provided the user-friendly open-source computational package, ASSISTER, for this framework. On the other hand, in the absence of rapid reversible binding, one can reduce a model by assuming that ‘highly reactive species’ are in their QSSs [58]. Interestingly, the reduced stochastic models were often different from the heuristically reduced models obtained with the deterministic QSSA. It would be interesting in future work to investigate when such discrepancies occur.

While the deterministic tQSSA (Eq. (3)) was used to approximate the stochastic QSSA for the number of reversibly binding species in this work, a simpler deterministic QSSA referred to as the “standard” QSSA (sQSSA) is more widely used to approximate the stochastic QSSA due to its simplicity [8–20, 27, 32]. For instance, the stochastic sQSSA for *C* in Eq. (1), which has the Michaelis-Menten type form

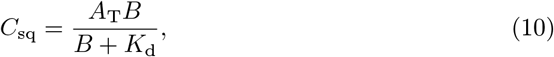

has been widely used as a propensity function for the Gillespie algorithm. Though this sQSSA has been widely used, it is less accurate than the stQSSA (Eq. (5)) [38, 39]. This is why many examples showing the inaccuracy of the stochastic sQSSA have been reported [33–40], whereas only one example showing the inaccuracy of the stQSSA has been reported [41]. Note that while Eq. (10) is different from the typical “Michaelis-Menten” equation, which uses the Michaelis-Menten constant instead of the dissociation constant (*K*_d_), they become nearly the same when the timescale of reversible binding is faster than the catalytic reaction. Importantly, our work also provides the validity condition for using the stochastic sQSSA (Eq. (10)). That is, when *B*_T_ + *K*_d_ ≫ *A*_T_, which is known as the low-enzyme concentration condition, *C*_sq_ ≈ *C*_tq_ [7], indicating that the Michaelis-Menten type sQSSA for the bounded species (Eq. (10)) can be used to reduce models containing the rapid reversible binding. Similarly, when *B*_T_ + *K*_d_ ≫ *A*_T_, 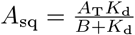 could also be used. This is consistent with the validity conditions for the stochastic sQSSA derived under the assumption of either a low fluctuation level [32] or a low copy number [40]. Furthermore, the “pre-factor” QSSA (pQSSA), which is more accurate than the sQSSA, has also been used for stochastic simulations [63, 64]. However, recent studies have shown that the stQSSA is more accurate than the stochastic pQSSA (see [38, 39] for details).

The accuracy of the stQSSA for the number of the unbound species depends on both the molar ratio between reversibly binding species and the tightness of their binding (Fig 1d). However, as the molar ratio typically varies in larger models containing reversible binding, practically, the accuracy is mainly determined by the tightness of binding. Specifically, for the relative error of the stQSSA to be less than 0.1, *A*_T_*K*_d_ (≈ the number of the unbound A) should be larger than 10 (Fig 5a dashed line). This *A*_T_*K*_d_ value-based criteria explains the controversy about the accuracy of the stQSSA in previous studies. That is, *A*_T_*K*_d_ was less than 10 in a previous study where the reduced model obtained by using the stQSSA was inaccurate [41]. On the other hand, *A*_T_*K*_d_ were much greater than 10 in all of the examples investigated in previous studies reporting the accuracy of the stQSSA [7, 33, 38, 39, 53, 54]. Furthermore, the stQSSA always accurately approximates the stochastic QSSA for the number of the bound species (Fig 1c). This explains why the stQSSA was accurate in previous studies where the stQSSA was used to approximate the number of enzyme-substrate complex [30–33].

In real biological systems, the validity condition of the stQSSA (*A*_T_*K*_d_ > 10) is not always guaranteed. Specifically, the range of *A*_T_*K*_d_ can span approximately from 10^−3^ to 10^10^ in human cells (Ω = 10^−15^ ~ 10^−14^*m*^3^) since the protein-protein dissociation constant 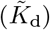 is 10*f M* ~ 1*μM* (i.e., 10^12^ ~ 10^20^*m*^−3^), and the numbers of molecules (*A*_T_) is 10^0^ ~ 10^4^ [43, 65]. Moreover, in smaller cells like budding yeast (Ω = 10^−17^ ~ 10^−16^*m*^3^) or E. Coli (Ω = 10^−19^ ~ 10^−18^*m*^3^) cells, the range of *A*_T_*K*_d_ can span from 10^−7^ to 10^8^, which contains the region in which the stQSSA can be extremely inaccurate. Accordingly, the slQSSA, which accurately approximates the stochastic QSSA when *A*_T_*K*_d_ is less than 10, is necessary. Specifically, the relative error of the slQSSA, unlike that of the stQSSA (Fig 5a and 5b), decreases as *A*_T_*K*_d_ decreases because the slQSSA relies on the assumption that the stationary distributions of the number of the unbound species (≈*A*_T_*K*_d_) are concentrated on the few lowest states. Taken together, by using the stQSSA and the slQSSA when the *A*_T_*K*_d_ value is greater and less than 10, respectively, one can always accurately simplify stochastic models containing rapid reversible binding reactions to accelerate simulation and also facilitate stochastic analysis (Fig 5f). This can be facilitated by the computational package, ASSISTER (Fig 6).

## Methods

### Exact bounds for the relative error of the stQSSA to the stochastic QSSA

In this section, we derive the exact upper and lower bounds for 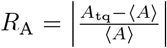 (Eq. (7)) where *A*_tq_ and ⟨*A*⟩ are the stQSSA and the stochastic QSSA for *A*, respectively. From the CME describing the reversible binding reaction (Eq. (1)), the following steady-state moment equation can be derived:

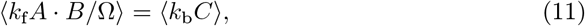

where ⟨·⟩ is the stationary expectation. Eq. (11) becomes ⟨*A* · (*B*_T_ − *A*_T_ + *A*)⟩ = *K*_d_⟨*A*_T_ − *A*⟩ by using the definitions *A*_T_ = *A* + *C*, *B*_T_ = *B* + *C*, and *K*_d_ = *k*_b_Ω*/k*_f_. Since *A*_T_ and *B*_T_ are invariant under the reversible binding reactions in Eq. (1), we obtain ⟨*A*^2^⟩ − (*A*_T_ − *B*_T_ − *K*_d_)⟨*A*⟩ − *A*_T_*K*_d_ = 0, and by using the relation ⟨*A*^2^⟩ = Var(*A*) + ⟨*A*⟩^2^, we get the following quadratic equation:

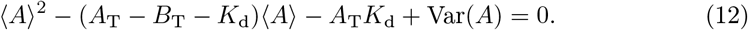

The non-negative root of this quadratic equation becomes ⟨*A*⟩:

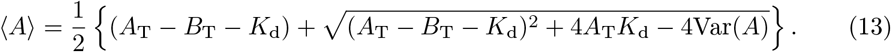

By subtracting Eq. (13) from Eq. (5), we get

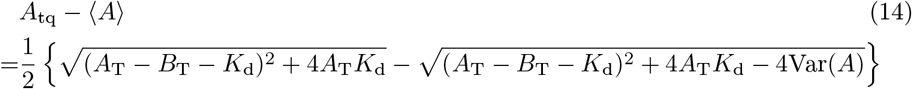

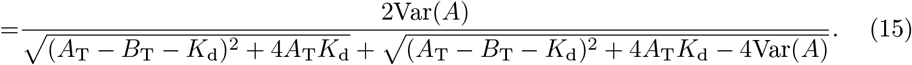

Since 0 ≤ (*A*_T_ − *B*_T_ − *K*_d_)^2^ + 4*A*_T_*K*_d_ − 4Var(*A*) ≤ (*A*_T_ − *B*_T_ − *K*_d_)^2^ + 4*A*_T_*K*_d_, we get the bounds for *A*_tq_ − ⟨*A*⟩ from Eq. (15):

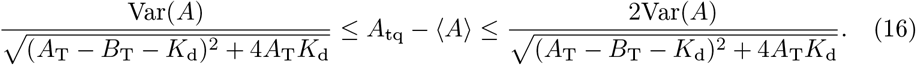

By dividing Eq. (16) by ⟨*A*⟩, we can get the bounds for the relative error, 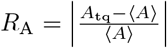 as follows:

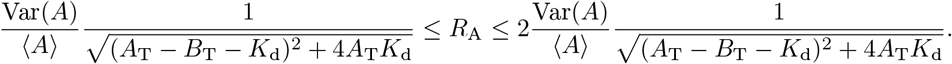

This can be re-expressed as *F*_A_*S*_A_ ≤ *R*_A_ ≤ 2*F*_A_*S*_A_ (Eq. (7)) because 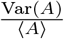 is the Fano factor of *A* (*F*_A_), and 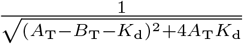 is the relative sensitivity of *A*_tq_, i.e., 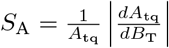.

The relative sensitivity, *S*_A_, attains the maximum value 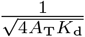 when the term in the square root of the denominator has the minimum value, i.e., *B*_T_ = *A*_T_ − *K*_d_ (Eq. (8)). In particular, *S*_A_ has a large maximum value when *K*_d_ ≪ 1 at *A*_T_ = *B*_T_ + *K*_d_ ≈ *B*_T_. On the other hand, if *A*_T_ ≪ *B*_T_, *S*_A_ ≈ 0 because the majority of A presents in the bound state regardless of *B*_T_ (i.e., 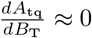). When *A*_T_ ≥ *B*_T_, 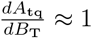 because as *B*_T_ decreases by one, approximately one A is released from the complex. In this case, if *A*_T_ ≫ *B*_T_, the majority of A are free and thus 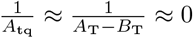, leading to *S*_A_ ≈ 0. However, if *A*_T_ ≈ *B*_T_, the majority of A is sequestered by B, *A*_tq_ ≈ 0, leading to *S*_A_ ≫ 1. When binding is weak (*K*_d_ ≫ 1), *S*_A_ ≈ 0 because the number of A, which is approximated by *A*_tq_, changes little as *B*_T_ changes (i.e., 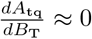). Taken together, *S*_A_ is large only when the binding reaction is tight (*K*_d_ ≪ 1) and the binding species are present with 1:1 molar ratio (*A*_T_ ≈ *B*_T_).

Since 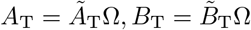, and 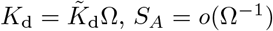, *S_A_* = *o*(Ω^−1^) if the concentrations remain constant. This implies that when the volume Ω goes to infinity (i.e., thermodynamic limit), *S_A_* and thus *R_A_* become zero (i.e., the stochastic QSSA becomes nearly identical to the deterministic QSSA (tQSSA)). On the other hand, as Ω goes to zero (i.e., the volume of the system gets smaller), *S_A_* goes to infinity.

### Derivation of the stochastic QSSA and the slQSSA

Here we derive the stochastic QSSA for *A* (⟨*A*⟩, Eq. (4)). Let *p*(*l*) be the probability that *A* = *l* at its stationary distribution (i.e., the probability that *A*(∞) = *l*). Then the following recurrence relation of *p*(*l*) can be obtained from the steady-state CME:

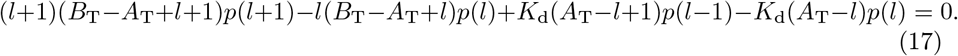

Let *A*_0_ = max{*A*_T_ − *B*_T_, 0}. Since *A*_0_ is the lowest state that *A* can reach, *p*(*l*) = 0 for *l < A*_0_. Then we can inductively prove that the following relation satisfies Eq. (17):

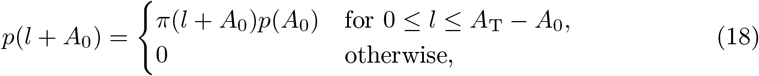

Where 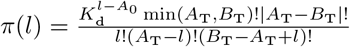. Then, because ∑*p*(*l*) = 1, 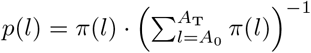 if *A*_0_ ≤ *l* ≤ *A*_T_, and *p*(*l*) = 0 otherwise by Eq. (18).

Therefore, we can obtain the stationary average number of A (Eq. (4)) as

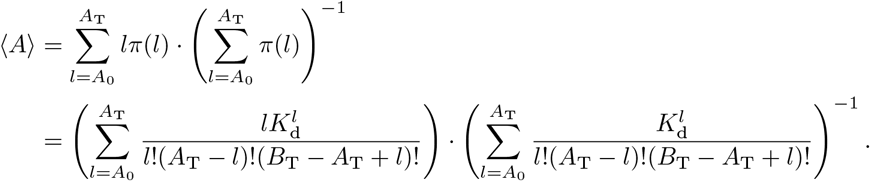

Next we derive the slQSSA, which is the approximation for Eq. (4). In the presence of tight binding, we can assume that the stationary distributions of *A* and *B* are concentrated on the states {0, 1} when *A*_T_ < *B*_T_ and *A*_T_ ≥ *B*_T_, respectively. Since when the distribution of *B* is concentrated on 0 and 1, the distribution of *A* is concentrated on *A*_T_ − *B*_T_ and *A*_T_ − *B*_T_ + 1, we can simply say that the distribution of A is concentrated on *A*_0_ and *A*_0_ + 1. Thus, by assuming that 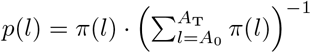 is approximately zero for *l* > *A*_0_ + 1 and 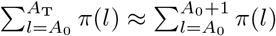, we can derive the two-state slQSSA for *A* (Eq. (9)) as follows:

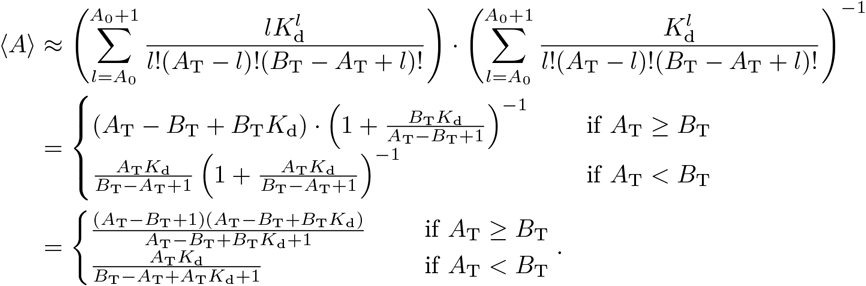

In general, for any integer *k* ≥ 2, we can derive the *k*-state slQSSA as

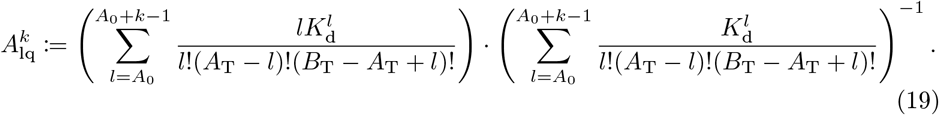

### Computational package for universally valid reduction of stochastic models containing rapid reversible binding reactions

We have developed a user-friendly computational package ASSISTER that contains three main codes implemented in MATLAB (Fig 6): `LQSSA, QSSA_Threshold`, and `Gillespie_Reduction. LQSSA` calculates the *L*-state slQSSA (Eq. (19)) for given *A*_T_, *B*_T_, *K*_d_, and *L*. `QSSA_Threshold` determines which of the stQSSA and the *L*-state slQSSA ensures a smaller error than a tolerance *ϵ* for a given *K*_d_ value. This allows the function `Gillespie_Reduction` to perform accurate stochastic simulations for any values of the parameters with the adaptive choice of the valid approximation method determined by using `QSSA_Threshold` (Fig 6). See S1 Appendix for details and the manual. ASSISTER can be found at https://github.com/Mathbiomed/ASSISTER.

## Supporting information

S1 Appendix. Supplementary Methods, Tables S1-S6, and Figs S1-S3. (PDF)

## Acknowledgments

We thank Krešimir Josić and John J. Tyson for valuable comments. This work was funded by NRF-2016 RICIB 3008468 (JKK), the Institute for Basic Science IBS-R029-C3 (JKK), and NRF-2019-Fostering Core Leaders of the Future Basic Science Program/Global Ph.D. Fellowship Program 2019H1A2A1075303 (HH).

## Author Contributions

All authors designed the study and performed mathematical analysis. YS performed the computation and all authors analyzed the computation results. YS and JKK wrote the draft and all authors revised the manuscript.

