## Appendix for "Universally valid reduction of multiscale stochastic biochemical systems using simple non-elementary propensities" (from another version of this preprint: version 1, 2021-04-09)

#### Supplementary Methods

##### Exact bound for the Fano factor of $A$

In this section, we show that the Fano factor of $A$ ($F_{\mathrm{A}}$) is always less than 1. We first define $\langle A^n|a,b\rangle$ as the stationary $n^{th}$ power moment of $A$ conditioned on $A_{\mathrm{T}} = a$ and $B_{\mathrm{T}} = b$. Then for any positive integer $a$ and $b$,

$$\begin{aligned} F_{\mathrm{A}} &= \frac{\langle A^2|a,b\rangle - \langle A|a,b\rangle^2}{\langle A|a,b\rangle} \\ &= \frac{\langle A^2|a,b\rangle}{\langle A|a,b\rangle} - \langle A|a,b\rangle. \end{aligned} \tag{S1}$$

When $a = 1$, $A$ can be only 0 and 1, so $\langle A^2|1,b\rangle = \langle A|1,b\rangle$, and thus
$F_{\mathrm{A}} = 1 - \langle A|1,b\rangle < 1$. Now we assume that $a \geq 2$. From Theorem 2 in [1], we have

$$\langle A^{(r)}|a,b\rangle = K_{\mathrm{d}}^r \frac{F_0(a-r,b)}{F_0(a,b)} \tag{S2}$$

for any $a \geq r$ and $b \geq 0$ where $A^{(r)} = \frac{A!}{(A-r)!}$. Here, $F_0$ is a generating function defined by

$$F_0(b_1,b_2) = \sum_{\substack{k_1,k_2,k_3\geq 0,\\ k_1+k_3=b_1,\\ k_2+k_3=b_2}} \frac{\lambda_1^{k_1}}{k_1!}\frac{\lambda_2^{k_2}}{k_2!}\frac{\lambda_3^{k_3}}{k_3!}$$

where $(\lambda_1, \lambda_2, \lambda_3)$ is any positive deterministic steady state of the reversible binding reaction [2]. Here, we set $(\lambda_1, \lambda_2, \lambda_3) = (K_{\mathrm{d}}, 1, 1)$. Then for $r = 1$,

$$\langle A|a,b\rangle = K_{\mathrm{d}} \frac{F_0(a-1,b)}{F_0(a,b)}, \tag{S3}$$

and for $r = 2$,

$$\langle A^2|a,b\rangle - \langle A|a,b\rangle = K_{\mathrm{d}}^2 \frac{F_0(a-2,b)}{F_0(a,b)}. \tag{S4}$$

We can rewrite Eq. (S4) as

$$
\begin{aligned}
\langle A^2|a,b\rangle &= K_{\mathrm{d}}^2 \frac{F_0(a-2,b)}{F_0(a,b)} + \langle A|a,b\rangle \\
&= K_{\mathrm{d}} \frac{F_0(a-2,b)}{F_0(a-1,b)} \cdot K_{\mathrm{d}} \frac{F_0(a-1,b)}{F_0(a,b)} + \langle A|a,b\rangle \\
&= \langle A|a-1,b\rangle \cdot \langle A|a,b\rangle + \langle A|a,b\rangle. \quad (\because \text{Eq. (S3)})
\end{aligned}
$$

Then by replacing $\langle A^2|a,b\rangle$ in Eq. (S1), we get

$$
F_{\mathrm{A}} = 1 + \langle A|a-1,b\rangle - \langle A|a,b\rangle. \tag{S5}
$$

Since $\langle A|a-1,b\rangle < \langle A|a,b\rangle$, we finally proved $F_{\mathrm{A}} < 1$.

While $\langle A|a-1,b\rangle < \langle A|a,b\rangle$ seems trivial, for the rigorousness, we provide the proof for this as follows. Since $F_{\mathrm{A}}$ is always positive from Eq. (S5),

$$
\langle A|a-1,b\rangle + 1 > \langle A|a,b\rangle, \tag{S6}
$$

for all $a \geq 2$ and $b > 0$. From the recurrence relation derived in the previous work [1], we have

$$
F_0(a,b) = \frac{K_{\mathrm{d}} + 1 + b - a}{a} F_0(a-1,b) + \frac{K_{\mathrm{d}}}{a} F_0(a-2,b),
$$

which can be re-expressed as

$$
\frac{F_0(a,b)}{F_0(a-1,b)} = \frac{K_{\mathrm{d}} + 1 + b - a}{a} + \frac{K_{\mathrm{d}}}{a} \frac{F_0(a-2,b)}{F_0(a-1,b)}.
$$

By multiplying $K_{\mathrm{d}}$ to both sides, we have

$$
\langle A|a,b\rangle^{-1} = K_{\mathrm{d}} \left( \frac{K_{\mathrm{d}} + 1 + b - a}{a} + \frac{1}{a} \langle A|a-1,b\rangle \right).
$$

Then we can show $\langle A|a-1,b\rangle < \langle A|a,b\rangle$, as follows:

$$
\begin{aligned}
\langle A|a,b\rangle &= K_{\mathrm{d}}^{-1} \left( \frac{K_{\mathrm{d}} + 1 + b - a}{a} + \frac{1}{a} \langle A|a-1,b\rangle \right)^{-1} \\
&> K_{\mathrm{d}}^{-1} \left( \frac{K_{\mathrm{d}} + 1 + b - a}{a} + \frac{1}{a} \left( \langle A|a-2,b\rangle + 1 \right) \right)^{-1} \quad (\because \text{Eq. (S6)}) \\
&> K_{\mathrm{d}}^{-1} \left( \frac{K_{\mathrm{d}} + 2 + b - a + \langle A|a-2,b\rangle}{a-1} \right)^{-1} \\
&= K_{\mathrm{d}}^{-1} \left( \frac{K_{\mathrm{d}} + 2 + b - a}{a-1} + \frac{1}{a-1} \langle A|a-2,b\rangle \right)^{-1} \\
&= \langle A|a-1,b\rangle.
\end{aligned}
$$

### Supplementary Tables

**Table S1. Propensity functions of the full model for the simple gene regulatory network (Fig 2a left)**

| Reactions | Propensity functions | Rate constants ($s^{-1}$) |
|---|---|---|
| D + P → D:P | $\frac{k_\mathrm{f}}{\Omega} DP$ | $k_\mathrm{f}/\Omega = 10^2, 10^2, 1, 10^2$ * |
| D:P → D + P | $k_\mathrm{b} D{:}P$ | $1, 1, 10, 1$ * |
| D → D + $M_\mathrm{R}$ | $k_\mathrm{R} D$ | $10^{-1}$ |
| D:P → D:P + $M_\mathrm{A}$ | $k_\mathrm{A} D{:}P$ | $10^{-3}$ |
| $M_\mathrm{R} \to \emptyset$ | $\tilde{k}_\mathrm{R} M_\mathrm{R}$ | $10^{-3}$ |
| $M_\mathrm{A} \to \emptyset$ | $\tilde{k}_\mathrm{A} M_\mathrm{A}$ | $10^{-3}$ |

* These four values are used for Figs 2b, 2c, 2d, and 5c, respectively.
[$D$, $P$, $D{:}P$, $M_\mathrm{R}$, $M_\mathrm{A}$] is initially [10, 10, 0, 0, 0], [15, 10, 0, 0, 0], [10, 10, 0, 0, 0], and [10, 10, 0, 0, 0] in Figs 2b, 2c, 2d, and 5c, respectively.

**Table S2. Propensity functions of the reduced models for the simple gene regulatory network obtained using the stQSSA (Fig 2a right) and the slQSSA (Fig 5c)**

| Reactions | Propensity functions | | Rate constants ($s^{-1}$) |
|---|---|---|---|
| | stQSSA | slQSSA | |
| $\emptyset \to M_\mathrm{R}$ | $k_\mathrm{R} D_\mathrm{tq}$ [1] | $k_\mathrm{R} D_\mathrm{lq}$ [3] | $10^{-1}$ |
| $\emptyset \to M_\mathrm{A}$ | $k_\mathrm{A} D{:}P_\mathrm{tq}$ [2] | $k_\mathrm{A} D{:}P_\mathrm{lq}$ [4] | $10^{-3}$ |
| $M_\mathrm{R} \to \emptyset$ | $\tilde{k}_\mathrm{R} M_\mathrm{R}$ | | $10^{-3}$ |
| $M_\mathrm{A} \to \emptyset$ | $\tilde{k}_\mathrm{A} M_\mathrm{A}$ | | $10^{-3}$ |

[1] $D_\mathrm{tq} = \{(D_\mathrm{T} - P_\mathrm{T} - K_\mathrm{d}) + \sqrt{(D_\mathrm{T} - P_\mathrm{T} - K_\mathrm{d})^2 + 4D_\mathrm{T}K_\mathrm{d}}\}/2$

[2] $D{:}P_{tq} = \{(D_\mathrm{T} + P_\mathrm{T} + K_\mathrm{d}) - \sqrt{(D_\mathrm{T} - P_\mathrm{T} - K_\mathrm{d})^2 + 4D_\mathrm{T}K_\mathrm{d}}\}/2$

[3] $D_\mathrm{lq} = \begin{cases} (D_\mathrm{T} - P_\mathrm{T} + 1)(D_\mathrm{T} - P_\mathrm{T} + P_\mathrm{T}K_\mathrm{d})(D_\mathrm{T} - P_\mathrm{T} + D_\mathrm{T}K_\mathrm{d} + 1) - 1 & \text{if } D_\mathrm{T} \geq P_\mathrm{T} \\ (D_\mathrm{T}K_\mathrm{d})(P_\mathrm{T} - D_\mathrm{T} + D_\mathrm{T}K_\mathrm{d} + 1)^{-1} & \text{if } D_\mathrm{T} < P_\mathrm{T} \end{cases}$

[4] $D{:}P_\mathrm{lq} = \begin{cases} D_\mathrm{T} - (D_\mathrm{T} - P_\mathrm{T} + 1)(D_\mathrm{T} - P_\mathrm{T} + P_\mathrm{T}K_\mathrm{d})(D_\mathrm{T} - P_\mathrm{T} + D_\mathrm{T}K_\mathrm{d} + 1)^{-1} & \text{if } D_\mathrm{T} \geq P_\mathrm{T} \\ D_\mathrm{T} - (D_\mathrm{T}K_\mathrm{d})(P_\mathrm{T} - D_\mathrm{T} + D_\mathrm{T}K_\mathrm{d} + 1)^{-1} & \text{if } D_\mathrm{T} < P_\mathrm{T} \end{cases}$

[$M_\mathrm{R}$, $M_\mathrm{A}$] is initally [0, 0] and $P_\mathrm{T} = 10$. $D_\mathrm{T} = 10, 15, 10, 10$ and $K_\mathrm{d} = 10^{-2}, 10^{-2}, 10, 10^{-2}$ in Figs 2b, 2c, 2d, and 5c, respectively.

**Table S3. Propensity functions of the full model for the biological oscillator (Fig 3a top)**

| Reactions | Propensity functions | Rate constants ($s^{-1}$) |
|---|---|---|
| A + R → A:R | $\frac{k_\mathrm{f}}{\Omega} AR$ | $k_\mathrm{f}/\Omega = 10^6$ |
| A:R → A + R | $k_\mathrm{b} A{:}R$ | $10^2$ |
| A → A + M | $\alpha_\mathrm{M} A$ | 20 |
| M → M + $\mathrm{P}_1$ | $\alpha_\mathrm{P} M$ | 1.5 |
| $\mathrm{P}_1$ → $\mathrm{P}_1$ + $\mathrm{P}_2$ | $\alpha_\mathrm{P} P_1$ | 1 |
| $\mathrm{P}_2$ → $\mathrm{P}_2$ + $\mathrm{P}_3$ | $\alpha_\mathrm{P} P_2$ | 1 |
| $\mathrm{P}_3$ → $\mathrm{P}_3$ + R | $\alpha_\mathrm{R} P_3$ | 1 |
| M → ∅ | $\beta_\mathrm{M} M$ | 1 |
| $\mathrm{P}_i$, $i = 1, 2, 3 \to \emptyset$ | $\beta_\mathrm{P} P_i$ | 1 |
| R → ∅ | $\beta_\mathrm{R} R$ | $0.02, 1, 0.02^*$ |
| A:R → A | $\beta_\mathrm{R} A{:}R$ | $0.02, 1, 0.02^*$ |

* These three values are used for Figs 3b, 3d, and 5d, respectively.
$[A,\ R,\ A{:}R,\ M,\ P_1,\ P_2,\ P_3]$ is initially [5, 5, 0, 0, 0, 0, 0].

**Table S4. Propensity functions of the reduced models for the biological oscillator obtained using the stQSSA (Fig 3a bottom) and the slQSSA (Fig 5d)**

| Reactions | Propensity functions | | Rate constants ($s^{-1}$) |
|---|---|---|---|
| | stQSSA | slQSSA | |
| ∅ → M | $\alpha_\mathrm{M} {A_\mathrm{tq}}^1$ | $\alpha_\mathrm{M} {A_\mathrm{lq}}^2$ | 20 |
| M → M + $\mathrm{P}_1$ | $\alpha_\mathrm{P} M$ | | 1.5 |
| $\mathrm{P}_1$ → $\mathrm{P}_1$ + $\mathrm{P}_2$ | $\alpha_\mathrm{P} P_1$ | | 1 |
| $\mathrm{P}_2$ → $\mathrm{P}_2$ + $\mathrm{P}_3$ | $\alpha_\mathrm{P} P_2$ | | 1 |
| $\mathrm{P}_3$ → $\mathrm{P}_3$ + $\mathrm{R_T}$ | $\alpha_\mathrm{R} P_3$ | | 1 |
| M → ∅ | $\beta_\mathrm{M} M$ | | 1 |
| P → ∅ | $\beta_\mathrm{P} P$ | | $0.02, 1, 0.02^*$ |
| $\mathrm{R_T}$ → ∅ | $\beta_\mathrm{R} R_\mathrm{T}$ | | $0.02, 1, 0.02^*$ |

[1] $A_\mathrm{tq} = \{(A_\mathrm{T} - R_\mathrm{T} - K_\mathrm{d}) + \sqrt{(A_\mathrm{T} - R_\mathrm{T} - K_\mathrm{d})^2 + 4A_\mathrm{T}K_\mathrm{d}}\}/2$

[2] $A_\mathrm{lq} = \begin{cases} (A_\mathrm{T} - R_\mathrm{T} + 1)(A_\mathrm{T} - R_\mathrm{T} + R_\mathrm{T}K_\mathrm{d})(A_\mathrm{T} - R_\mathrm{T} + A_\mathrm{T}K_\mathrm{d} + 1)^{-1} & \text{if } A_\mathrm{T} \geq R_\mathrm{T} \\ (A_\mathrm{T}K_\mathrm{d})(R_\mathrm{T} - A_\mathrm{T} + A_\mathrm{T}K_\mathrm{d} + 1)^{-1} & \text{if } A_\mathrm{T} < R_\mathrm{T} \end{cases}$

* These three values are used for Figs 3b, 3d, and 5d respectively.
$[R_\mathrm{T},\ M,\ P_1,\ P_2,\ P_3]$ is initially [5, 0, 0, 0, 0], $A_\mathrm{T} = 5$, and $K_\mathrm{d} = 10^{-4}$.

**Table S5. Propensity functions of the full model for the bistable switch (Fig 4a top) [3, 4]**

| Reaction | Propensity function | Rate constants ($s^{-1}$) |
|---|---|---|
| M + B → M:B | $\frac{k_\mathrm{f}}{\Omega}MB$ | $k_\mathrm{f}/\Omega = 10^4, 10, 10^{4*}$ |
| M:B → M + B | $k_\mathrm{b}M{:}B$ | $10^1, 10^2, 10^{1*}$ |
| M → P | $k_\mathrm{p}M$ | 0.6 |
| M:B → M + P | $k_\mathrm{p}M{:}B$ | 0.6 |
| P → M | $\left(k_1 + (k_2 - k_1)\frac{M}{K_\mathrm{D}+M}\right)DP$ | $k_1 = 0.05$, $k_2 = 1.1$ |

* These three values are used for Figs 4b, 4d, and 5e, respectively.
$[M,\ B,\ M{:}B,\ P]$ is initially [0, 10, 0, 30], $D = 1$, $K_\mathrm{D} = 0.1$.

**Table S6. Propensity functions of the reduced models for the bistable switch obtained using the stQSSA (Fig 4a bottom) and the slQSSA (Fig 5e) [3, 4]**

| Reactions | Propensity functions | | Rate constants ($s^{-1}$) |
|---|---|---|---|
| | stQSSA | slQSSA | |
| $\mathrm{M_T}$ → P | $k_\mathrm{p}M_\mathrm{T}$ | | 0.6 |
| P → $\mathrm{M_T}$ | $\left(k_1 + (k_2 - k_1)\frac{M_\mathrm{tq}}{K_\mathrm{D}+M_\mathrm{tq}}\right)DP^1$ | $\left(k_1 + (k_2 - k_1)\frac{M_\mathrm{lq}}{K_\mathrm{D}+M_\mathrm{lq}}\right)DP^2$ | $k_1 = 0.05$, $k_2 = 1.1$ |

[1] $M_\mathrm{tq} = \{(M_\mathrm{T} - B_\mathrm{T} - K_\mathrm{d}) + \sqrt{(M_\mathrm{T} - B_\mathrm{T} - K_\mathrm{d})^2 + 4M_\mathrm{T}K_\mathrm{d}}\}/2$

[2] $M_\mathrm{lq} = \begin{cases} (M_\mathrm{T} - B_\mathrm{T} + 1)(M_\mathrm{T} - B_\mathrm{T} + B_\mathrm{T}K_\mathrm{d})(M_\mathrm{T} - B_\mathrm{T} + M_\mathrm{T}K_\mathrm{d} + 1)^{-1} & \text{if } M_\mathrm{T} \geq B_\mathrm{T} \\ (M_\mathrm{T}K_\mathrm{d})(B_\mathrm{T} - M_\mathrm{T} + M_\mathrm{T}K_\mathrm{d} + 1)^{-1} & \text{if } M_\mathrm{T} < B_\mathrm{T} \end{cases}$

$[M_\mathrm{T},\ P]$ is initially [0, 30], $D = 1$, $K_\mathrm{D} = 0.1$, and $B_\mathrm{T} = 10$. $K_\mathrm{d} = 10^{-3}, 10, 10^{-3}$ in Figs 4b, 4d, and 5e, respectively.

#### Supplementary Figures

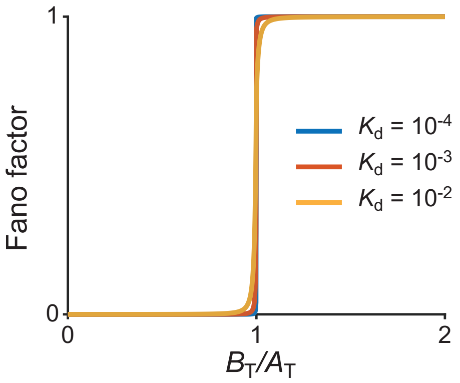


**Fig S1. The Fano factor of $A$ ($F_A$) is between 0 and 1.** As $K_d$ decreases, $F_A$ becomes closer to the step function, which is 0 when $A_T > B_T$ and 1 when $A_T \le B_T$. Since the relative error of the stQSSA for $A$ to the stochastic QSSA ($R_A$) satisfies $F_A S_A \le R_A \le 2F_A S_A$ (Eq. (7) in the main text), $R_A$ is almost 0 when $B_T < A_T$ while $R_A$ is between $S_A$ and $2S_A$ when $B_T \ge A_T$. Therefore, when A tightly binds with B and $B_T \ge A_T$, $R_A$ mainly depends on $S_A$ (Fig 1d and 1e). Here, $A_T = 100$. See Supplementary Methods for the proof that $F_A$ is always between 0 and 1.

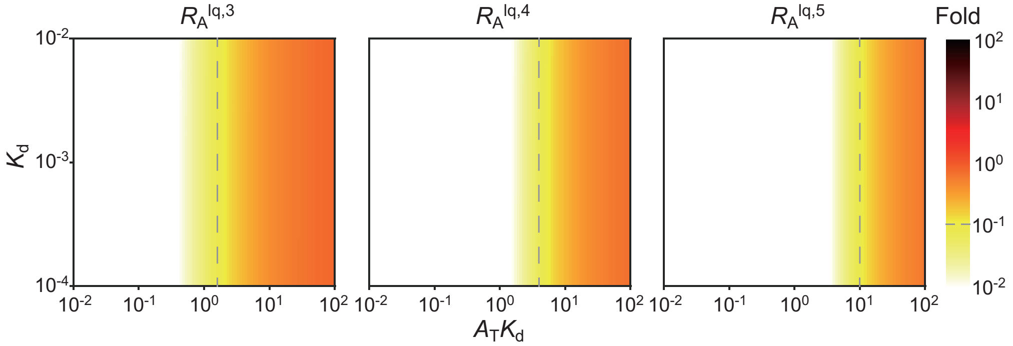


**Fig S2. slQSSAs with more states approximate the stochastic QSSA more accurately.** The heat maps of the relative errors ($R_A^{lq,k} = \left|\frac{A_{lq}^k - \langle A \rangle}{\langle A \rangle}\right|$) when the $k$-state slQSSA ($A_{lq}^k$) approximates the stochastic QSSA for $A$ ($\langle A \rangle$) for $k = 3, 4, 5$. See Eq. (18) in the main text for a detailed definition of $A_{lq}^k$. Color in the heat maps represents the maximum value of $R_A^{lq,k}$ for each $A_T K_d$ and $K_d$ when $B_T$ varies. As more states are used (i.e., $k$ increases), $A_{lq}^k$ accurately approximates $\langle A \rangle$ in the wider regions. Specifically, when $A_T K_d$ is less than 2, 5, and 10 (i.e., left side of the dashed lines in the heat maps), $R_A^{lq,3}$, $R_A^{lq,4}$ and $R_A^{lq,5}$ are less than 0.1, respectively. This indicates that one can accurately approximate $\langle A \rangle$ with relative error of less than 0.1 regardless of the values of $A_T$, $B_T$, and $K_d$ by adaptively using both $A_{tq}$ and $A_{lq}^5$ because the relative error of $A_{tq}$ is less than 0.1 when $A_T K_d > 10$.
